# Electromembrane extraction and mass spectrometry for liver organoid drug metabolism studies

**DOI:** 10.1101/2020.05.15.095174

**Authors:** Frøydis Sved Skottvoll, Frederik Hansen, Sean Harrison, Ida Sneis Boger, Ago Mrsa, Magnus Saed Restan, Matthias Stein, Elsa Lundanes, Stig Pedersen-Bjergaard, Aleksandra Aizenshtadt, Stefan Krauss, Gareth Sullivan, Inger Lise Bogen, Steven Ray Wilson

## Abstract

Liver organoids are emerging tools for precision drug development and toxicity screening. We demonstrate that electromembrane extraction (EME) based on electrophoresis across an oil membrane is suited for segregating selected organoid-derived drug metabolites prior to mass spectrometry (MS)-based measurements. EME, allowed drugs and drug metabolites to be separated from cell medium components (albumin, etc.) that could interfere with subsequent measurements. Multi-well EME (Parallel-EME) holding 100 μL solutions allowed for simple and repeatable monitoring of heroin phase I metabolism kinetics. Organoid Parallel-EME extracts were compatible with ultrahigh-performance liquid chromatography (UHPLC) used to separate the analytes prior to detection. Taken together, liver organoids are well-matched with EME followed by MS-based measurements.

## Introduction

The process of drug development is known to be time consuming and bear financial uncertainties^1,2^. It is estimated that from 5,000-10,000 new molecular entities, only one new drug will enter the market^3^. The advancement of this one drug from concept to market takes approximately 15 years and a cost over $ 1 billion, as well as the use of human resources, research skills, and technological expertise^3^. As the majority of drug candidates are rejected late in the process and during clinical trials^3^, one approach to reducing the assets put into the drug development may be to reject possible drug candidates early in the development process, i.e. during preclinical testing. This may be done by developing or utilizing in vitro models that adequately recapitulate the human in vivo response.

Organoids are three-dimensional tissue models derived from primary tissues, embryonic stem cells or induced pluripotent stem cells (iPSC)^4–6^. These “mini” organs are emerging tools for studying human development and disease, serving as alternatives to cell cultures and animal models in drug development^7,8^. A wide variety of organoids are being developed and studied, e.g. brain, heart, tumor tissue and liver^9–12^. Liver organoids can be valuable models for studying drug metabolism and toxicity^13^ (**Figure 1A**), perhaps even in a personalized fashion, as organoids can be derived from the cells of a patient^14,15^.

**Figure 1.**
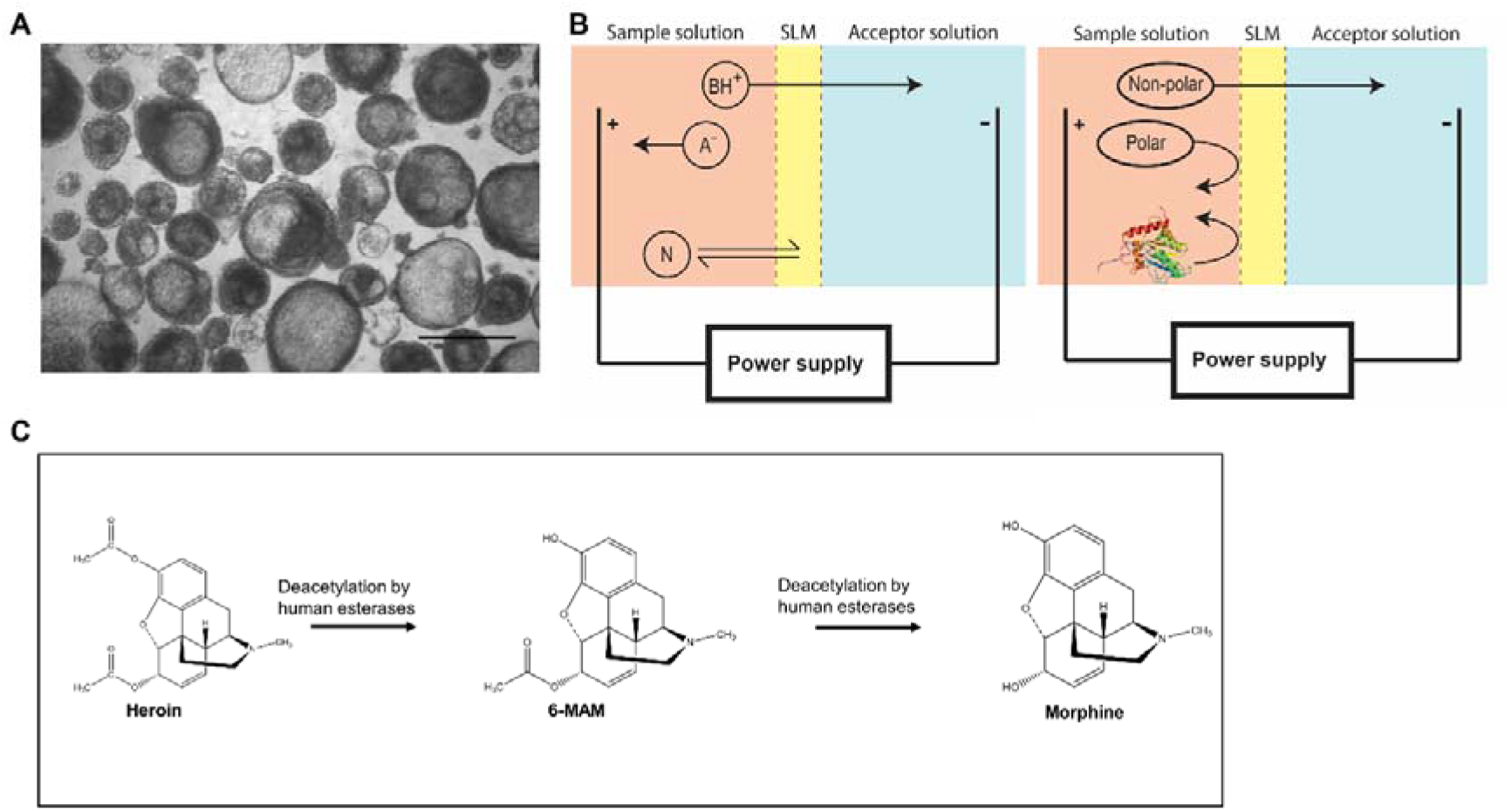
(**A**) Light microscope picture of iPSC derived liver organoids used in this study, scale bar 500 μm. (**B**) EME principle. Charged analytes migrate from the sample solution across the SLM and into the acceptor solution. Extraction selectivity is obtained by voltage polarity and partitioning into and through the SLM. Polar molecules and macromolecules are effectively discriminated from extraction by the hydrophobic SLM. (**C**) Illustration of well-documented liver phase I metabolism of heroin undergoing deacetylation to 6-MAM and morphine by human esterases (e.g. human liver carboxylesterase 1 and 2, hCE1 and hCE2).

Drug metabolism is a significant determinant of drug clearance and an indirect determinant of the clinical efficacy and toxicity of drugs^16^. Thus, the mapping of the biotransformation pathway of drugs is crucial in the early part of the drug development process^17^. Clinical studies of xenobiotics in humans are subjected to constraints concerning ethical aspects. Several in vitro model systems have been developed to recapitulate human functions from the molecular level to the cellular, tissue, organ, or whole organism level. The most commonly used in vitro models for drug metabolism studies include subcellular fractions e.g. human liver microsomes (HLMs), S9 fractions and human hepatocytes. However, current in vitro models have some disadvantages. For example HLMs do not represent a complete course of metabolism as they lack soluble phase II enzymes^16^. Additionally, higher biotransformation rates are obtained in HLMs compared to humans, most likely because of the enriched enzyme concentrations and the absence of competing enzymes^17^. Also, animal models can have shortcomings, and have frequently been shown to lead to wrong predictions of drug interaction and toxicity in humans^18^.

For both in vitro and in vivo models, drug metabolism studies are very often performed utilizing liquid chromatography-mass spectrometry (LC-MS). Essentially, the mass spectrometer (MS) can measure the drugs and their metabolites with a high degree of selectivity. Prior to MS measurements, the compounds in the sample are separated by the LC system, allowing for increased sensitivity and selectivity.

There are few studies utilizing LC-MS for drug metabolism measurements of organoids^19–21^. To the authors knowledge, there are currently no studies dedicated to demonstrating the potential of drug metabolism studies with liver organoids and LC-MS^22^. The key focus of this study is to show the potential of using liver organoids and LC-MS measurements as a methodology for drug metabolism studies. To ensure an efficient combination of organoids, LC-MS and drug metabolism, several challenges must be addressed. The amounts of organoids can (depending on the production method) be quite limited per sample, requiring efficient sample preparation prior to analysis. It is also highly desirable that drug metabolism studies with organoids can be upscaled, which is difficult to combine with more standard sample preparation approaches which include centrifugation steps and manual pipetting (**Figure S-1A**). In addition, liver organoids are grown in complex medium (e.g. can contain 10 % fetal bovine serum) requiring a thorough sample clean-up prior to LC-MS analysis. For extracting drugs, and the metabolites produced by organoids, we have applied electromembrane extraction (EME, **Figure 1B** and **Figure S-1B**). In EME, an oil immobilized in the pores of a porous membrane (supported liquid membrane, SLM) is used to extract analytes from a cell medium (donor solution) to a protein free MS compatible acceptor solution. For the process, both aqueous compartments are pH-adjusted to facilitate analyte ionization, and voltage is applied across the SLM. EME is therefore essentially an electrophoretic migration of ionized analytes across an oil membrane^23,24^. Extraction selectivity is determined by both the partitioning of analytes into the SLM, and the polarity and magnitude of the applied voltage. High clean-up efficiency of target analytes can thus be achieved, and EME is highly successful separating small-molecule drug substances from biological matrix substances, including salts, lipids, phospholipids, proteins, and blood cells^24,25^. Such clean-up is highly important prior to liquid chromatography-mass spectrometry to avoid ion suppression or enhancement. EME has recently advanced to the 96-well plate format^26–28^ (Parallel-EME), and chip systems^29,30^. Considering its documented traits regarding simple sample clean-up, we focus on using EME for organoids, which can be costly and limited in availability.

As a model system to show the potential of the methodology, we study the phase I metabolism of heroin to 6-monoacetylmorphine (6-MAM) and morphine (**Figure 1C**), as heroin liver metabolism is highly established, both with regards to the metabolizing enzymes^31–33^ (e.g. human liver carboxylesterase 1 and 2, hCE1 and hCE2), and the resulting metabolites. With the here presented experiments, we have shown proof of concept that liver organoids are EME compatible, and evaluate the advantages and challenges of Parallel-EME/organoid/MS-based analysis for drug metabolism.

## Experimental Section

### Chemicals and Solutions

2-Nitrophenyl octyl ether (NPOE), 2-nitrophenyl pentyl ether (NPPE), bis (2-ethylhexyl) hydrogen phosphite (DEHPi), bis(2-ethylhexyl) phosphate (DEHP), sodium hydroxide, ammonium formate (>99%), formic acid (FA, reagent grade 95%), L-ascorbic acid-2 phosphate (AAP) were purchased from Sigma Aldrich (St. Louis, MO, USA). LC-MS grade water and acetonitrile (ACN) was purchased from VWR (Radnor, PA, US). Chromasolv methanol (LC-MS grade) was from Honeywell Riedel-de Haën (Seelze, Germany). Heroin HCl, 6-MAM HCl and morphine were purchased from Lipomed AG (Arlesheim, Switzerland). Heroin-d9, 6-MAM-d6 and morphine-d3 were purchased from Cerilliant (Austin, TX, USA). Unless otherwise stated, the water used was type 1 water purified by a Milli-Q® water purification system from Merck Millipore (Billerica, MA, USA).

The 5 mM and 10 mM ammonium formate buffer (*w/v*) was made by dissolving ammonium formate in LC-MS grade water followed by pH adjustment by the addition of FA to pH 3.1. A freshly made stock solution of 1 mM heroin HCl in 0.9% NaCl was made prior to each organoid experiment (stored at 4 °C), and was also used to prepare heroin calibration solutions. A stock solution of 6-MAM and morphine was prepared in 5 mM ammonium formate buffer pH 3.1 at a concentration of 50 μM each and stored at 4 °C. Two stock solutions of the internal standards heroin-d9, 6-MAM-d6 and morphine-d3 were prepared in 5 mM ammonium formate buffer pH 3.1 with analyte concentration of 1.5 μM each and 3 μM each, respectively, and stored at 4 °C.

### Liver organoid differentiation from induced pluripotent stem cells

The iPSC cell line HPSI0114i-vabj_3 (Wellcome Trust Sanger Institute, Cambridgeshire, UK) was differentiated toward liver organoids using media from protocol by Ang et al.^34^. Briefly, the HPSI0114i-vabj_3 iPSC line was differentiated toward definitive endoderm in Iscove's Modified Dulbecco's Medium/F12 medium (Thermo Fisher Scientific, Waltham, MA, USA) containing 3 μM CHIR99021 (STEMCELL Technologies, Vancouver, Canada), 50 nM PI-103 from Bio-Techne Ltd. (Abingdon, United Kingdom) and 100 ng/mL activin A (PeproTech, Cranburdy, NJ, USA) for one day and 100 ng/mL activin A for 2 more days. The definitive endoderm cells were subsequently treated with 1 μM A8301 (Bio-Techne Ltd.), 10 ng/mL FGF2 (PeproTech), 30 ng/mL BMP4 (PeproTech), and 2 μM all-trans retinoic acid (Sigma Aldrich) for one day, then with 10 ng/mL FGF2, 30 ng/mL BMP4, 1 μM forskolin (PeproTech), 1 μM Wnt-C59 (Bio-Techne Ltd.) for 2 more days and with 10 ng/mL FGF2, 30 ng/mL BMP4, 1 μM forskolin for another day. At day 8 cells were detached and aggregated in the U bottom microwells in the presence of 20 ng/mL HGF (PeproTech), 10 ng/mL oncostatin M (OSM, PeproTech), 0.1 μM dexamethasone (Bio-Techne Ltd.), 1 μM forskolin, 10 μg/mL human recombinant insulin (Sigma Aldrich), 100 μM AAP. After formation of organoids at day 10 they were transferred into low attachment plates and cultured for another 10 days as free-floating organoids in William's E media (Thermo Fisher Scientific), supplemented with 10 ng/mL HGF and 10 ng/mL OSM, 10 μg/mL insulin, 100 μM AAP, 0.1 μM dexamethasone, 1 μM forskolin and 10 μM DAPT (Bio-Techne Ltd.). The iPSC line AG27^35–38^ was differentiated using a small molecule driven protocol that aims to sequentially mimic in vivo liver development, resulting in hepatocyte containing liver organoids as described by Harrison et al.^39^.

### Liver organoid heroin incubation

Prior to heroin incubation with organoids, 1 mM heroin was diluted in the respective cell medium and sterilized by filtration using a 0.22 μm Millex-GV syringe filter (Merck Millipore). After 20 days differentiation, from 20 to 60 organoids per well were treated with 10 or 50 μM heroin in cell medium for 1, 3, 6, and 24 hours respectively (n=3), in separate Nunc flat-bottom 96-well microplates (Thermo Fisher Scientific). Metabolism was stopped by adding FA to a final concentration of 0.11 M, and the plates were frozen at −80 °C. In parallel, cell medium free from organoids (n=3) were used as drug degradation control samples.

### Parallel electromembrane extraction setup

Prior to the extraction, 50 μL of the heroin-exposed liver organoid samples (containing 0.11 M FA) was added to 40 μL water and 10 μL of the 1.5 μM or 3 μM internal standard solution. The samples were then loaded into the wells of an in-house built 96-well stainless-steel plate (**Figure 2A**), previously described by Restan et al.^28^. A volume of 3 μL DEHP/NPOE (10/90, *w/w*) was immobilized into the membrane pores (0.45 μm pore size) of a 96-well MultiScreen-IP polyvinylidene fluoride (PVDF) filter plate from Merck Millipore (**Figure 2B**). The steel and filter plates were subsequently clamped together and 100 μL of 10 mM ammonium formate pH 3.1 was loaded into each well of the filter plate, and thus constituting the acceptor solution. The filter plate was used to house the acceptor solution because the geometry of the steel plate wells provided better convection of the sample solution in this configuration, which improved the extraction kinetics. A conductive in-house built aluminum lid with 96 electrode rods (**Figure 2C**) was placed onto the filter plate, and the whole construct (**Figure 2D**) was placed on a Vibramax 100 Heidolph shaking board (Kellheim, Germany). The steel plate holding the organoid solution was connected to the anode of an external power supply (model ES 0300e0.45, Delta Elektronika BV, Zierikzee, The Netherlands), while the aluminum electrode lid was connected to the cathode (**Figure 2E**). Simultaneous extraction of all samples was performed for 15 min at 900 rpm agitation, with 30 V applied for the first two minutes and 50 V applied for the remaining extraction duration. The stepped voltage was used to ensure that the extraction current was kept below 50 μA per well, which was considered a safe limit for robust operation^40^.

**Figure 2.**
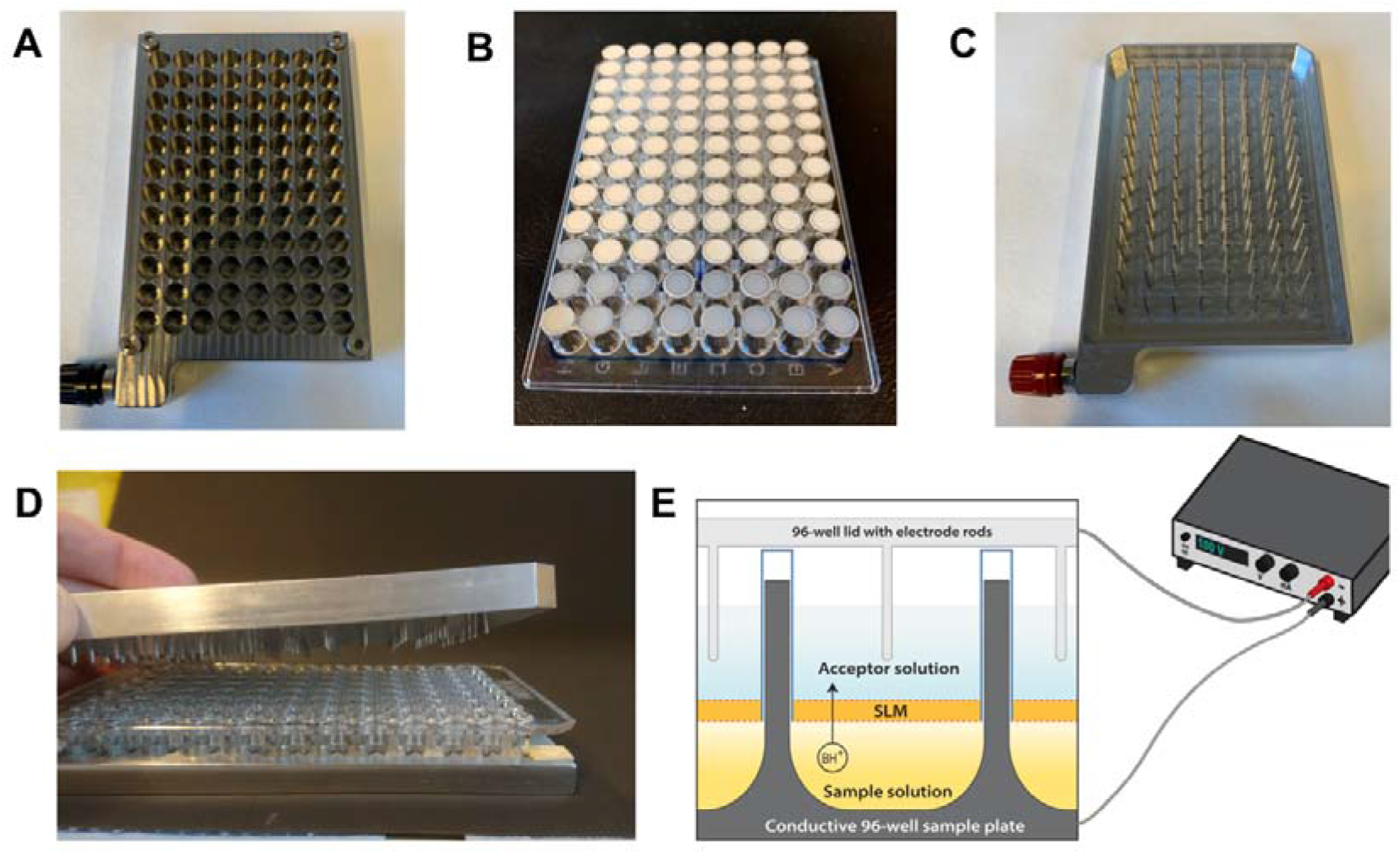
The experimental setup of 96 well Parallel-EME. (**A**) The 96 well sample reservoir plate constituting the donor solution. (**B**) 96 well filter plate, constituting the acceptor solution. (**C**) The aluminum lid with 96 electrode rods. (**D**) All plates clamped together. (**E**) Illustration of the extraction setup of Parallel-EME coupled to the external power supply.

### Ultra high-performance liquid chromatography-mass spectrometry (UHPLC-MS)

Determination of heroin, 6-MAM and morphine was performed using UHPLC-MS based on a previously described method^41^. The sample extracts were diluted x10 with 5 mM ammonium formate pH 3.1 and analyzed using an Acquity™ UHPLC pump coupled to a Xevo TQ (triple quadrupole) MS with an electrospray ionization (ESI) interface, all from Waters (Milford, MA, USA). Separation was achieved using the Acquity UPLC® HSS T3 C18 column (2.1 × 100 mm, 1.8 μm particles). Solvent A consisted of 10 mM ammonium formate buffer pH 3.1 and solvent B consisted of methanol. The sample injection volume was set to 7.5 μL, and the gradient elution was carried out at a flow rate of 0.5 mL/min at 65 °C using the following gradient profile: from 0–0.5 min; 100% solvent A, 0.5–2.7 min; 0-10% solvent B, 2.7–3.3 min; 10%–20% solvent B, 3.3–4.6 min; 20%–80% solvent B, 4.6–4.61 min; 80%–100% solvent B, 4.61-6.60 min; 100% solvent B, 6.60–6.61 min; 100%–0% solvent B, 6.61–7.50 min; 100% solvent A. The capillary voltage was 3 kV, source temperature 150 °C, desolvation temperature 500 °C and cone gas flow 990 L/h. Detection was performed in positive mode using multiple reaction monitoring (MRM) with MS/MS transitions (MS/MS transition 1 being the quantifier and MS/MS transition 2 the qualifier) and collision energies for heroin (*m/z* 370> 268 at 30 eV and *m/z* 370> 211 at 38 eV), 6-MAM (*m/z* 328> 165 at 42 eV and *m/z* 328> 211 at 30 eV), morphine (*m/z* 286> 201 at 24 eV and *m/z* 286> 165 at 42 eV), heroin-d9 (*m/z* 379> 272 at 30 eV), 6-MAM-d6 (*m/z* 334> 165 at 42 eV) and morphine-d3 (289> 165 at 30 eV). Data was acquired and processed using MassLynx 4.1 software (Waters).

### Nano liquid chromatography mass spectrometry (nanoLC-MS)

The nanoLC-MS setup consisted of a TSQ Quantiva, triple quadrupole MS, the nanoFlex ESI ion source and the EASY-nLC 1000 or 1200 pump equipped with an autosampler, all from Thermo Fisher. Acclaim PepMap™ 100 C18 (3 μm particle size) pre- (75 μm inner diameter, ID, and 20 mm length) and analytical (75 μm ID x 50 mm) columns from Thermo Fisher Scientific were used for the chromatographic separation. In-house made^42^ analytical columns were packed with 3 μm Atlantis T3 particles (Waters) or 2.6 μm Accucore particles (Thermo Fisher Scientific) in fused silica capillaries of 75 μm ID from Polymicro Technologies (Phoenix, AZ, USA). The analytical column was coupled to a 40 mm stainless steel emitter (20 μm ID) purchased from Thermo Fisher. The extracted organoid samples (AG27 iPSC derived) were further diluted x10^3^ in 5 mM of ammonium formate pH 3.1 buffer, and the injection volume was set to 2 μL. The nanoLC pump was equipped with two solvent compartments (A and B), where A contained 0.1% FA in LC-MS grade water (*v/v*) and B contained 0.1% FA in LC-MS grade water and ACN (10/90, *v/v*). The gradient elution was carried out with 3-50% B in 8 min with a constant flow rate of 500 nL/min. The spray voltage was set to 2.2 kV and the ion transfer tube temperature was set to 310 °C. Detection was performed in positive mode using MRM with MS/MS transitions and collision energies for heroin (*m/z* 370> 268 at 38 eV and 370> 211 at 41 eV), 6-MAM (*m/z* 328> 165 at 48 eV and 328> 211 at 36 eV), morphine (*m/z* 286> 181 at 48 eV and 286> 165 at 51 eV), heroin-d9 (*m/z* 379> 272 at 38 eV and 379> 211 at 41 eV), 6-MAM-d6 (*m/z* 334> 211 at 35 eV and 334> 165 at 48 eV) and morphine-d3 (*m/z* 289> 181 at 48 eV and 289> 165 at 51 eV).

For a one-column setup, the pump outlet was coupled to an external six-port valve from Valco Instruments Company Inc (VICI®, Houston, TX, USA) equipped with a 75 μm ID x 11 cm fused silica injection loop (500 nL), a nut with a syringe sleeve and a 75 μm ID x 10 cm fused silica capillary waste outlet. The flow outlet from the 6-port valve was coupled to a stainless-steel tee-piece (VICI®) through a 20 μm x 40 cm fused silica capillary from Polymicro Technologies using stainless steel nuts and vespel/graphite ferrules (VICI®). The analytical column inlet was coupled to the stainless-steel tee piece, also coupled to a plug through a 550 mm nanoViper (75 μm ID, Thermo Fisher). A 500 μL syringe (51mm) from Hamilton (Reno, Nevada, USA) was used to load the samples. Xcalibur™ version 2.2 was used to obtain chromatograms and mass spectra (Thermo Fisher).

### Protein profiling by nano liquid chromatography mass spectrometry

Acetone precipitated AG27 iPSC derived liver organoid protein samples were subjected to SDS-PAGE gel electrophoresis, and the gel lanes were sliced into five sample fractions and digested with trypsin as previously described^43^. The peptide solutions were desalted using OMIX C18-micro solid phase extraction (SPE) columns (Agilent, Santa Clara, CA, USA). A Q-Exactive mass spectrometer (Thermo Fisher Scientific) equipped with a nanoFlex nanospray ion source was used for the nanoLC-MS analyses, coupled to an EASY-nLC 1000 pump (Thermo Fisher). Peptide separation was achieved using Acclaim PepMap 100 pre- (20 mm) and separation columns (250 mm) of 75 μm inner diameter and 3 μm particles (Thermo Fisher). Solvent A was 0.1% FA in LC-MS grade water (*v/v*), and solvent B was 0.1% FA in LC-MS grade water and ACN (5/95, *v/v*). Peptides were separated using a 180 minutes long gradient ranging from 3-15% solvent B (after optimization with pre-digested HeLa samples from Thermo Fisher). The mass spectrometer was run in positive mode with full MS (*m/z* = 400-2000) and data dependent tandem mass spectrometry (ddMS2) with top N set to be 10 ions. Raw files were processed and database searches performed with Proteome Discoverer 2.3 (Thermo Fisher Scientific), using MASCOT version 2.4 to search the SwissProt database (human, 20 431 entries). Proteins were identified with the following settings; peptide identification with a false discovery rate (FDR) threshold of ≤ 0.01, protein identification with a FDR threshold of ≤ 0.01 (strict) and ≤0.05 (relaxed) and digestion by trypsin with at most one missed cleavage. Dynamic modification was set to be oxidation and acetyl (N-term), static modification was set to be carbamidomethyl. Information on the elution profile and fragment match spectrum of each of the identified peptides for hCES1 (accession number P23141), hCES2 (also called cocaine esterase, accession number O00748) and UDP-glucuronosyltransferase 2B7 (accession number P16662) were obtained and verified by comparison with the raw file.

### Calculation of Recovery

Recovery measurements were performed using capillary electrophoresis with ultraviolet spectroscopy detection (CE-UV) (See supplementary for experimental description) with an initial analyte concentration of 5 μM. The recovery (%) was calculated using the following formula:

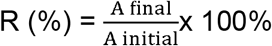

Where A_final_ and A_initial_ are the area of analyte collected in the acceptor solution and the area of the analyte originally present in the sample.

## Results and Discussion

In this study, several analytical approaches were evaluated for liver organoid drug measurements. With the future objective of advancing to online analyses, EME was assessed in 96-well format (Parallel-EME) for the high-throughput clean-up of analytes from the organoid cell medium, a method previously shown to enable selective and fast extraction from complex matrices (and also on-chip)^44^. A conventional UHPLC-MS method used for clinical routine analyses was applied to explore heroin metabolizing properties of the Parallel-EME extracted liver organoids. To get an understanding of the heroin metabolizing liver enzymes present in the organoids, an untargeted proteomic case study using nanoLC-MS was undertaken. Lastly, two analytical approaches more suitable for online action, limited samples, and increased sensitivity were evaluated: CE which is widely established for rapid on-chip separations ^45–47^, and nanoLC-MS allowing for high sensitivity measurements^48^.

### Parallel electromembrane extraction optimization for heroin and metabolites

To evaluate the potential of MS for analysis of liver organoids, heroin was chosen as a model substance, due to its familiar phase I metabolism to 6-MAM and morphine in the liver. Although morphine extraction with EME has previously been performed^49–51^, the extraction of heroin and 6-MAM with EME has to our knowledge not previously been performed. Therefore, Parallel-EME conditions focusing on these three compounds were initially assessed. The experimental conditions (**Figure 3**) were selected based on previous experience and literature reports^49,52,53^. Due to the difference in polarity of the analytes, > 30% recovery and < 15% RSD were set as the acceptance criteria of extraction performance. Best recovery and repeatability for analytes in both standard solutions and spiked cell medium samples were obtained using an Parallel-EME system comprising 10% (*w/w*) DEHP/NPOE as SLM, an extraction time of 15 min, and an extraction voltage of 50 V. From cell medium, these conditions gave recoveries of 76% (heroin), 82% (6-MAM), and 36% (morphine) and RSD < 10%, which was considered acceptable for the current application. The extraction method was therefore not optimized any further. With these parameters, the average extraction current was < 50 μA per well throughout the extraction. For increasing accuracy, correction for non-exhaustive extractions was done by spiking samples with isotopically labelled internal standards prior to extraction.

**Figure 3.**
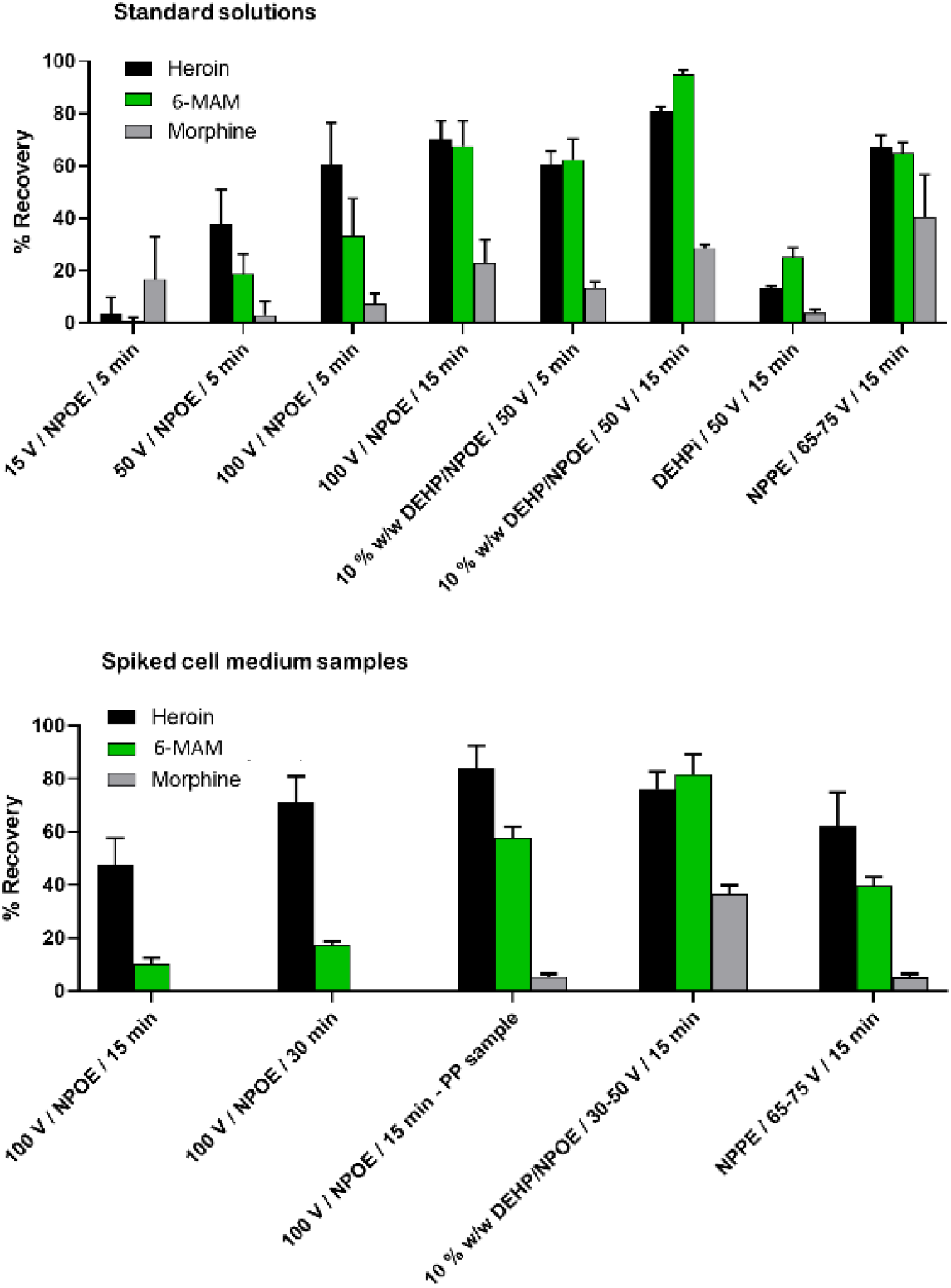
Analyte recovery (%) of Parallel-EME under varying conditions (SLM composition, extraction voltage and extraction time), with 5 μM standard solutions and spiked cell medium samples using CE-UV for quantitation.

### Parallel electromembrane extraction of liver organoid heroin metabolites

Samples containing 20 and 60 liver organoids per well were exposed to 10 μM heroin for 1, 3, 6 and 24 hours. With the exception of 6-MAM and heroin at time point 24 hours, the sample to sample repeatability was 0.4%-25% with the two organoid iPSC sources (**Figure 4A-B)**. Heroin levels decreased with time to 6-MAM (both enzymatic and non-enzymatic), and with subsequent enzymatic metabolism to morphine, adding to the confirmation that the liver organoids had traits related to human livers. Similar heroin metabolism kinetics was also observed for liver organoids derived from hepatocytes from one patient case (see **Figure S-2**). However, the kinetics were (expectedly) substantially slower than that observed with e.g. high enzyme-availability microsomes and S9-fraction^17,54^, see **Figure S-3**; Although Parallel-EME and MS are compatible with phase I metabolism monitoring, we were not able to observe phase II metabolites morphine-3-glucuronide (M3G) and morphine-6-glucuronide (M6G). Traces of these metabolites could however be observed when employing more manual, centrifugation-based sample preparation (**Figure S-4**). A key reason is a weakness of EME, that highly polar compounds have low recovery; this can in many cases be fine-tuned^53,55^.

**Figure 4.**
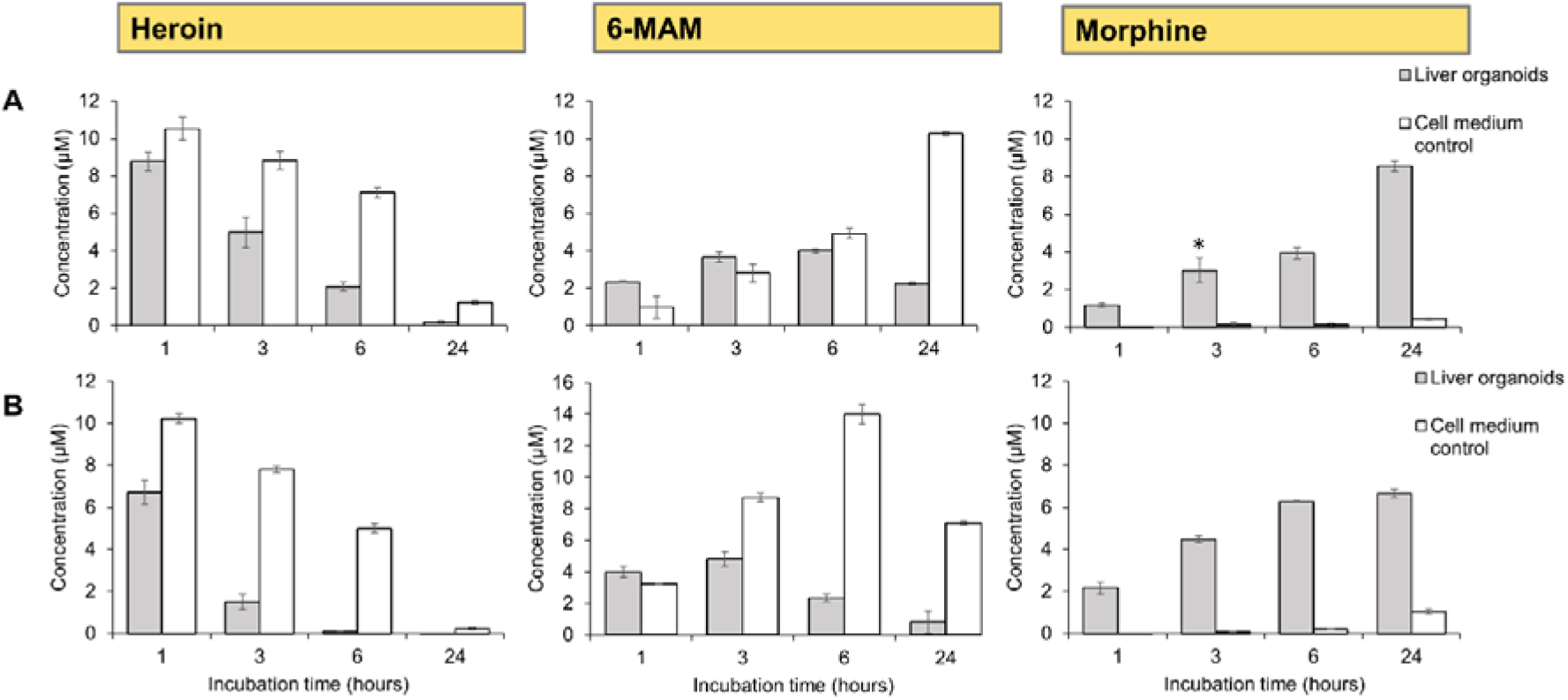
Concentration of heroin and metabolites in a study of liver organoid drug metabolism using Parallel-EME and UHPLC-MS, after incubation of liver organoids differentiated from the iPSC cell lines **(A)** AG27 (60 organoids) and **(B)** HPSI0114i-vabj_3 (20 organoids) in 10 μM heroin for 1, 3, 6- and 24 hours. In parallel, cell medium free from organoids were used as drug degradation control samples. Each bar represents the mean (± SD) of triplicate samples. One of the three replicates of time point 6 hours liver organoids (HPSI0114i-vabj_3) was discarded. The asterisk indicates the removal of one data point due to poor internal standard signal.

To complement the observations of the liver organoids enzymatic heroin metabolizing properties, a case study using MS-based untargeted proteomics was undertaken. We could identify the presence of proteotypic peptides (FDR ≤ 1%) related to the key liver enzymes^56–60^ hCES1 (9 peptides identified) and hCES2 (4 peptides identified) in the organoids differentiated from the iPSC cell line AG27 (**Figure 5A-C**, see also **Table S-1** for peptide overview). Also, one peptide was identified related to one of the heroin phase II metabolism enzymes^33,57^, UDP-glucuronosyltransferase 2B7 (**Table S-1**).

**Figure 5.**
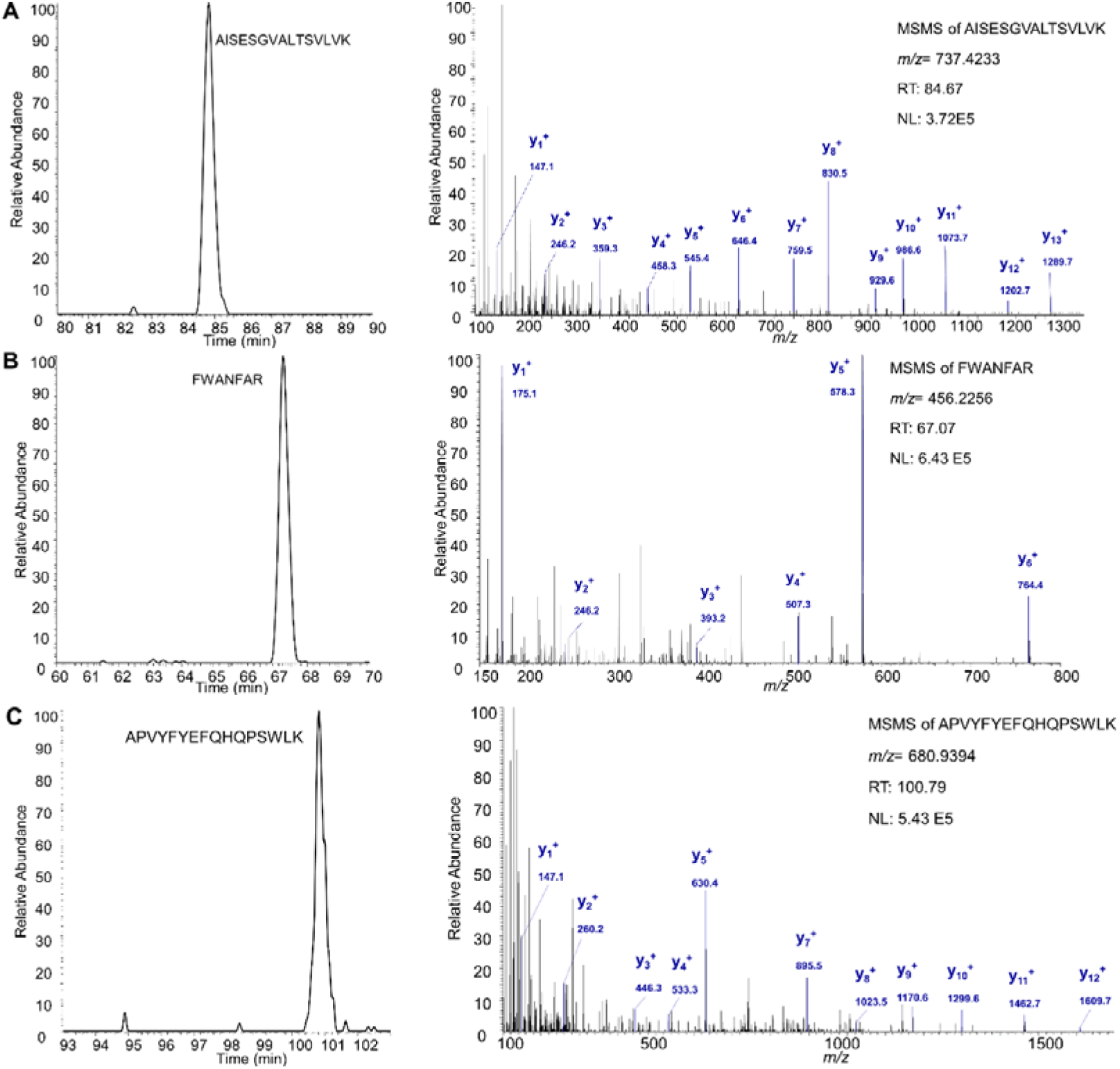
Total ion chromatogram of identified peptides (left) and the respective peptide fragmentation spectrum (right) of enzymes related to heroin liver phase I metabolism. (**A**) The peptide AISESGVALTSVLVK (*m/z* 737.42) from hCES1, identified at charge +2. (**B**) The peptide FWANFAR (*m/z* 456.23) from hCES1, identified at charge +2. (**C**) The peptide APVYFYEFQHQPSWLK (*m/z* 680.94) from hCES2, identified at charge +3.

### Organoid EME extracts compatibility with various separation techniques

The organoid EME extracts were analyzed using UHPLC-MS instrumentation, which provided high resolution separations within 5 min (**Figure S-5**). We have also investigated other separation approaches that can be compatible with small samples and online action. Capillary electrophoresis, perhaps the most “chip-ready” of the techniques investigated, was capable of fast separations of organoid extracts (separation within 2.5 min) and low sample consumption (injection volume equivalent to 107 nL), with these initial experiments demonstrated with simple UV detection (**Figure S-6**). However, organoid incubation in 50 μM heroin was needed to achieve detection with CE-UV, and thus no further quantification of the analytes could be performed.

The limit of quantification (LOQ) for UHPLC-MS measurements in this study was 1 nM (7 μL injection volume). NanoLC, a sensitive approach that has been mostly associated with proteomics in recent years, was seen to provide 0.95 pM detection (1 μL injection volume) for some small molecule analytes such as heroin (results not shown). The organoid extracts analyzed with nanoLC-MS could thus be 1 000 times more diluted compared to that of UHPLC-MS analysis, without compromising on chromatographic performance or sensitivity for the more hydrophobic analytes heroin and 6-MAM (**Figure 6A**). However, poor performance was associated with nanoLC-MS analysis of morphine, the most polar of the metabolites observed; the chromatographic peak was completely absent in the chromatograms of the organoid extracts (**Figure 6A**), and was sporadically very deformed or absent in that of standard solutions. This was the case for large volume injection, both using on-column injection and using an SPE column. We also examined in-house packed nano reversed phase (RP) LC columns which were more compatible with highly aqueous mobile phases (Accucore and Atlantis T3), but poor peak shape and breakthrough/poor retention time repeatability were still issues. Various parameters were tested, e.g. sample loading time and maximum sample loading pressure (of the Thermo nano pumps). To illustrate these effects, see **Figure 6B**, which shows that several loading times were suited for 6-MAM and heroin using on-column injection, but none were suited for morphine.

**Figure 6.**
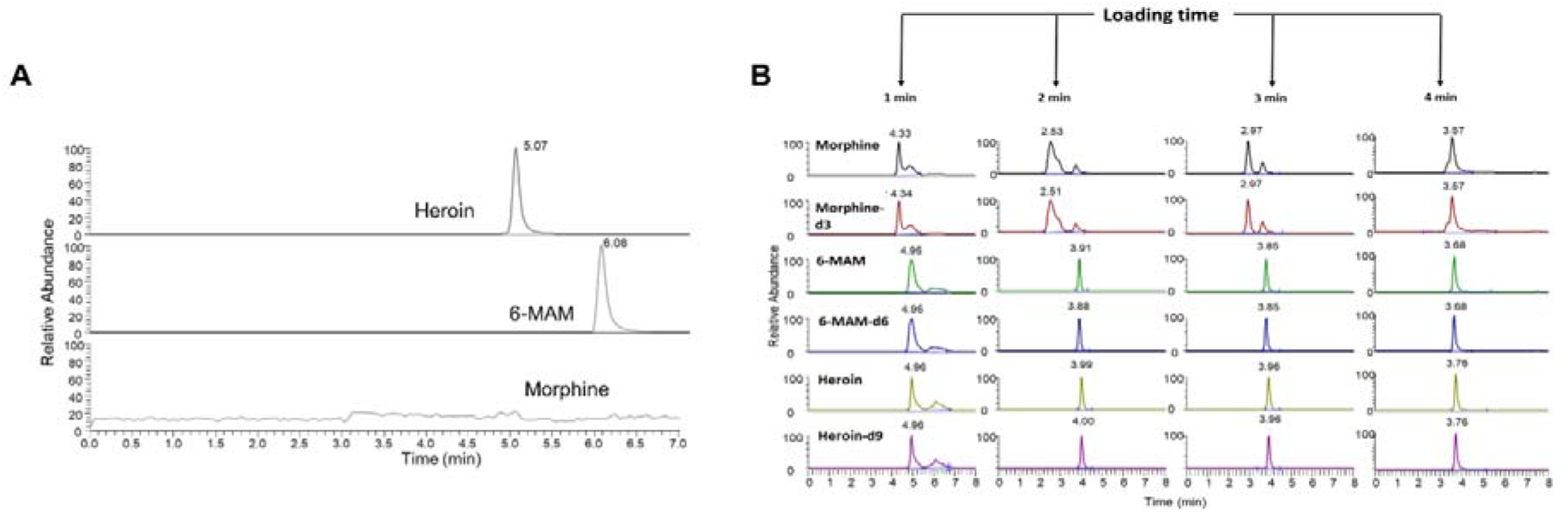
(**A**) MRM chromatograms of heroin, 6-MAM, and morphine in the extracted liver organoid sample treated with 10 μM heroin for 1 hours (AG27). The sample was analyzed using a two-column setup with Acclaim PepMap columns, and injection volume of 2 μL. (**B**) MRM-chromatograms of a 375 nM standard solution containing heroin, morphine, 6-MAM, and their corresponding internal standards, analyzed using the one-column setup equipped with an Acclaim PepMap commercially packed analytical column with different on-column loading times (1, 2, 3 and 4 min), and injection volume of 500 nL.

## Conclusions

Liver organoids and LC-MS measurements is a promising concept for drug metabolism studies, here demonstrated for heroin phase I metabolism. This concept can be well suited for drug metabolism studies of other drugs, and direct measurements of drug metabolism could also provide valuable insight when optimizing organoid development protocols. A proteomic case study using nanoLC-MS identified proteotypic peptides from heroin metabolizing enzymes, complementing the observations of the liver organoids enzymatic heroin metabolizing properties. EME-MS showed to be a promising combination for liver organoid based analysis of drug metabolism. EME in 96-well format (Parallel-EME) was used to extract heroin and metabolites from various organoids in a complex medium, followed by UHPLC-MS measurements. In addition, the chromatographic performance was not perturbed by the initial complex matrix (analyte retention time repeatability with a maximum RSD of 0.07%), suggesting that Parallel-EME was a suited basis for organoid derived sample preparation. It is reasonable to assume that the approach can also be applicable to other organoid variants, e.g. kidney and heart. Parallel-EME was indeed an approach that allowed multiple samples to be simply handled, more so than standard approaches to related tissues (centrifugations, several sample pipetting steps), which can allow higher throughput in larger-scale studies. We are currently developing 96-well plates made of conductive polymers, which we believe will be suited for both cell studies and EME; this will reduce yet another step of sample handling. One disadvantage that needs to be addressed is the difficulty in extracting very polar metabolites with EME, and further optimizations will therefore continue.

Following this proof-of-concept study, we will continue to explore iterations of the here presented EME-configuration with the aim of further increasing sensitivity while retaining robustness and scalability; a natural next step will be nanoliter-scale online EME-LC-MS of organoid derived samples. Related systems have been demonstrated with microsomes^30^, but those systems require larger separation columns, and are arguably not suited for trace samples. Due to challenges with nanoLC, we will instead likely investigate the use of capillary LC or microbore LC, as a compromise between sensitivity and robustness.

## Acknowledgements

This work was supported by the Research Council of Norway through its Centre of Excellence scheme, project number 262613. The work was also supported by the Olav Thon Foundation. Financial support from UiO:Life Science is also gratefully acknowledged. Technical assistance by Elisabeth Nerem (Department of Forensic Sciences, Oslo University Hospital, Oslo, Norway) was greatly appreciated.

