## Supporting Information for "Electromembrane extraction and mass spectrometry for liver organoid drug metabolism studies"

**Table of contents**

**Figure S-1:** S Schematic overview of the workflow of drug studies performed with (**A**) a standard sample preparation approach using centrifugation and several steps of manual pipetting prior to analysis, and (**B**) parallel electromembrane extraction (Pa-EME) with fewer steps of manual pipetting and sample handling prior to analysis.

**Figure S-2:** Liver organoid drug metabolism using Pa-EME and UHPLC-MS, after incubation of patient derived liver organoids differentiated from primary human hepatocytes (20 organoids) in 10 µM heroin for 1, 3, 6- and 24 hours. In parallel, cell medium free from organoids were used as drug degradation control samples. Each bar represents the mean (± SD) of triplicate samples.

**Figure S-3:** The concentration (µM) of heroin (0.1 µM final heroin concentration in incubation solution) and its phase I metabolites 6-MAM and morphine in (**A**) 0.01 mg microsomes and (**B**) 0.1 mg/mL S9-fraction after centrifugation based sample preparation. Separation and detection was performed using UHPLC-MS. Data points represent the mean (± SD) of triplicate samples, and the solid lines represent model prediction.

**Figure S-4:** Liver organoid drug phase I and II metabolism measuring the concentration (µM) of heroin, 6-MAM, morphine, morphine-3-glucuronide (M3G) and morphine-6-glucuronide (M6G) with the use of centrifugation based sample preparation and UHPLC-MS. Liver organoids differentiated from the iPSC cell line HPSI0114i-vabj_3 (20 organoids) were incubated in 10 µM heroin for 0 and 24 hours. Each bar represent the mean (± SD) of triplicate samples.

**Table S-1:** Identified peptides of the enzymes hCES1 (accession number P23141), hCES2 (accession number O00748) and UDP-glucuronosyltransferase 2B7 (accession number P16662) related to heroin liver metabolism, with annotated sequence, quality q-value, the total number (#) of protein groups identified with the peptide, number (#) of peptide spectral matches (PSMs), peptide position in the protein, number (#) of missed cleavages of the identified peptide. * The peptide is also identified in putative inactive carboxylesterase 4 (CES1P1, accession number Q9UKY3).

**Figure S-5:** Chromatogram showing the separation of the analytes M3G, M6G, morphine, 6-MAM and heroin with the use of UHPLC-MS.

**Figure S-6:** Electropherogram of liver organoid drug phase I metabolism using Pa-EME and simple capillary electrophoresis with ultraviolet spectroscopy detection (CE-UV), after incubation with liver organoids differentiated from the iPSC cell lines AG27 (60 organoids) in 50 µM heroin for 6 hours.

**Additional experimental procedures:** Heroin metabolism assay in S9-fractions and microsomes, sample preparation using centrifugation, capillary electrophoresis with ultraviolet spectroscopy detection.

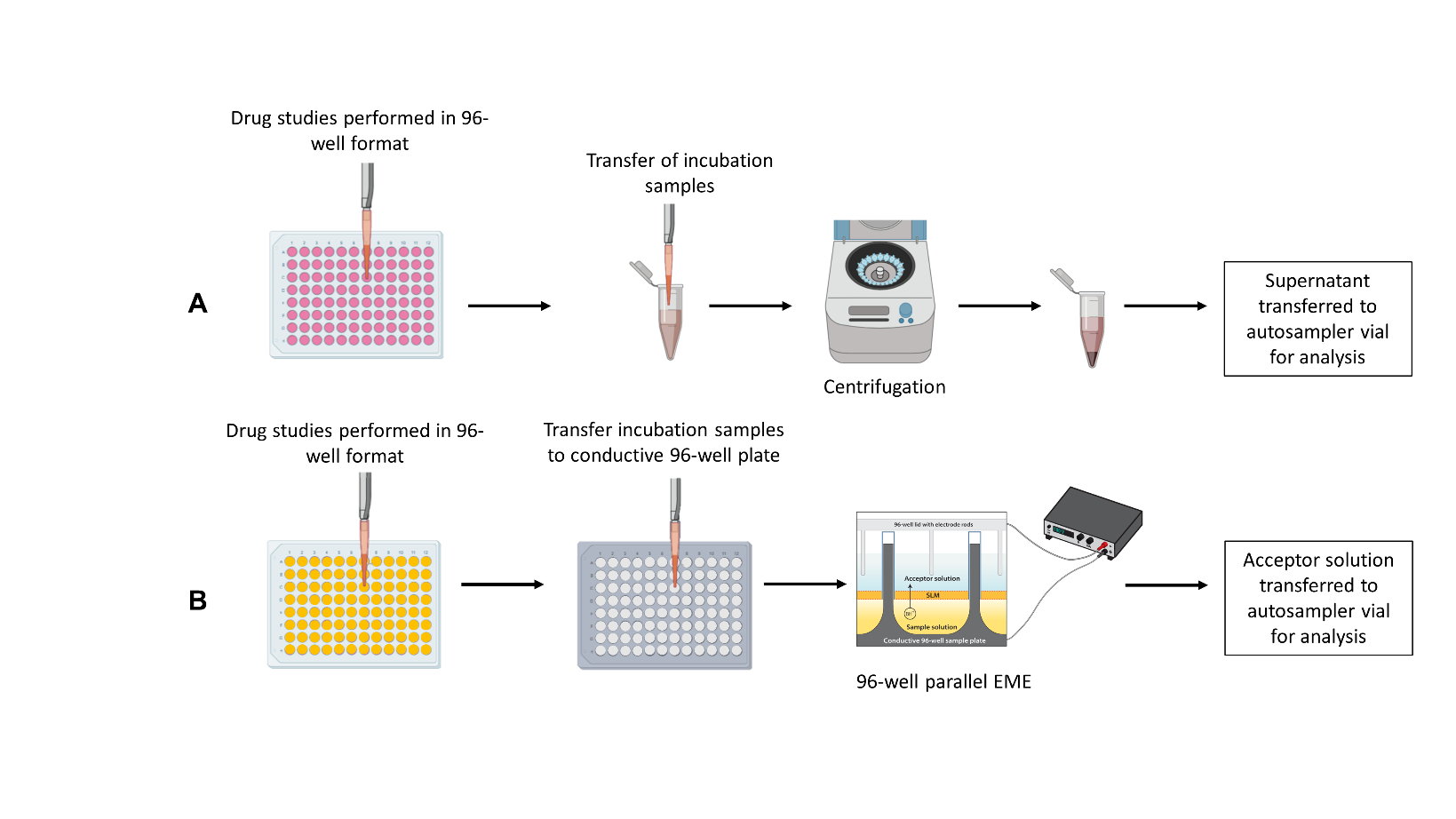

**Figure S-1.** Schematic overview of the workflow of drug studies performed with (**A**) a standard sample preparation approach using centrifugation and several steps of manual pipetting prior to analysis, and (**B**) parallel electromembrane extraction (Pa-EME) with fewer steps of manual pipetting and sample handling prior to analysis.

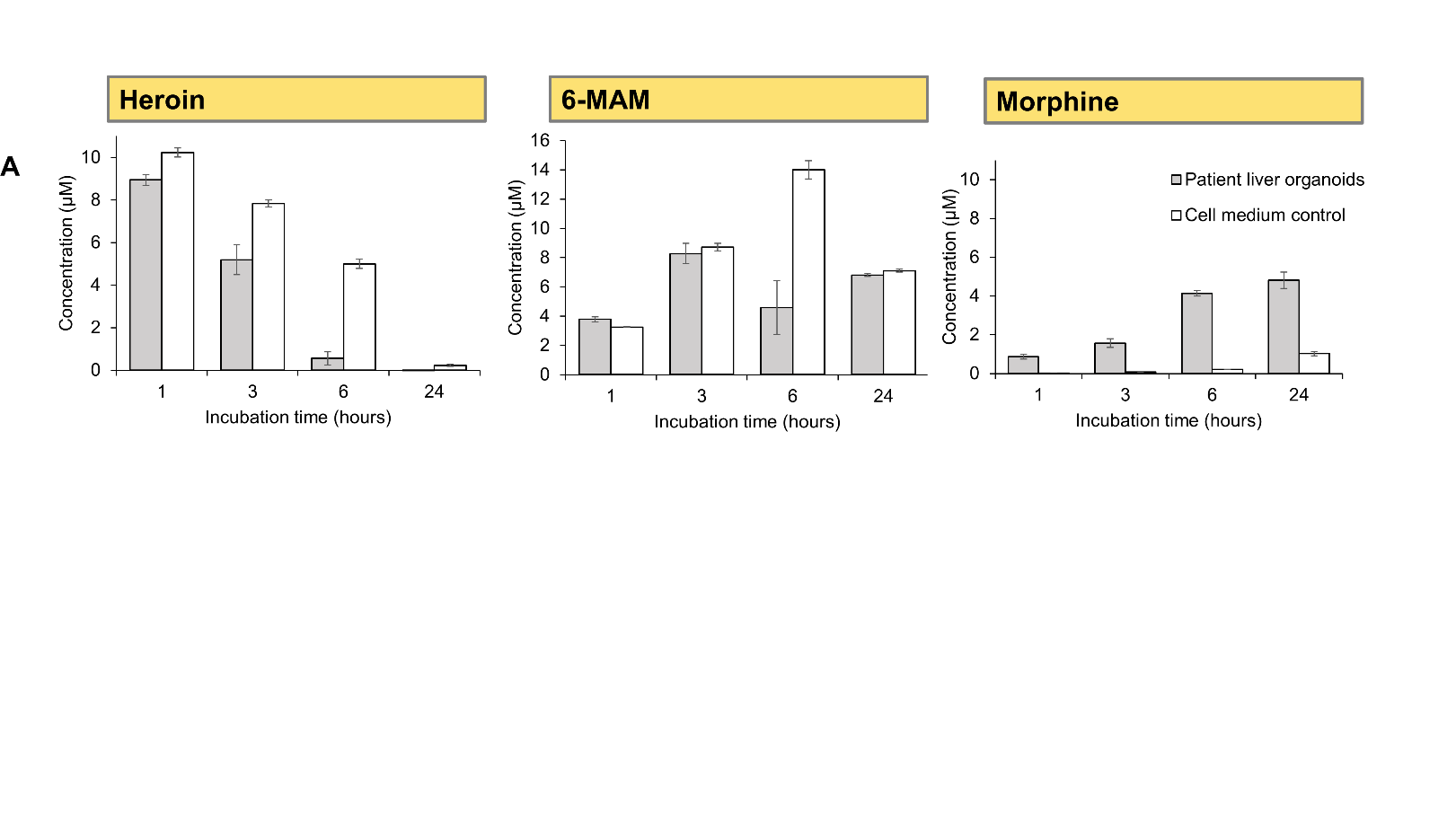

**Figure S-2.** Liver organoid drug metabolism using Pa-EME and UHPLC-MS, after incubation of patient derived liver organoids differentiated from primary human hepatocytes (20 organoids) in 10 µM heroin for 1, 3, 6- and 24 hours. In parallel, cell medium free from organoids were used as drug degradation control samples. Each bar represents the mean (± SD) of triplicate samples.

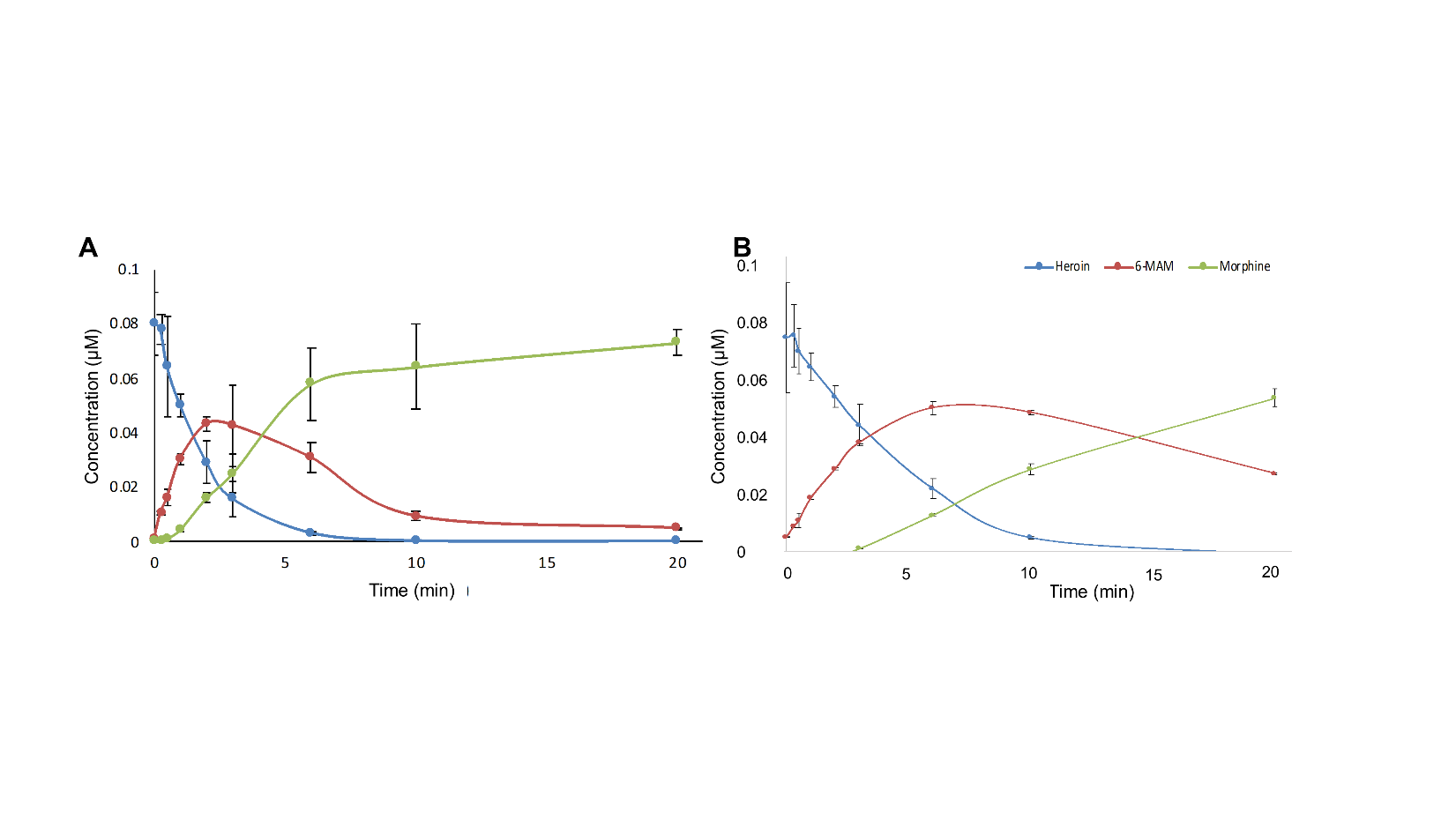

**Figure S-3.** The concentration (µM) of heroin (0.1 µM final heroin concentration in incubation solution) and its phase I metabolites 6-MAM and morphine in (**A**) 0.01 mg microsomes and (**B**) 0.1 mg/mL S9-fraction after centrifugation based sample preparation. Separation and detection was performed using UHPLC-MS. Data points represent the mean (± SD) of triplicate samples, and the solid lines represent model prediction.

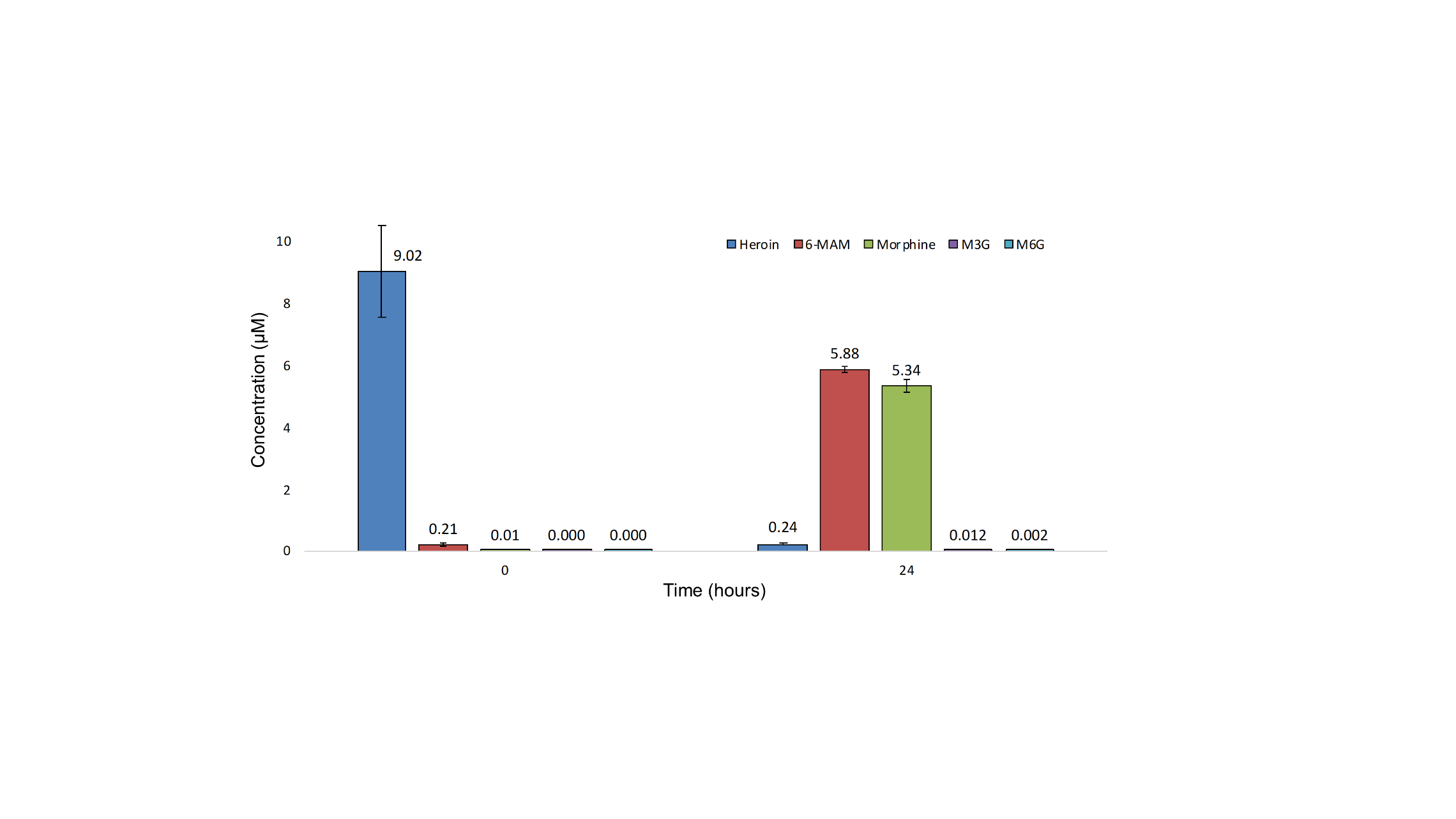

**Figure S-4**. Liver organoid drug phase I and II metabolism measuring the concentration (µM) of heroin, 6-MAM, morphine, morphine-3-glucuronide (M3G) and morphine-6-glucuronide (M6G) with the use of centrifugation based sample preparation and UHPLC-MS. Liver organoids differentiated from the iPSC cell line HPSI0114i-vabj_3 (20 organoids) were incubated in 10 µM heroin for 0 and 24 hours. Each bar represent the mean (± SD) of triplicate samples.

**Table S-1.**Identified peptides of the enzymes hCES1 (accession number P23141), hCES2 (accession number O00748) and UDP-glucuronosyltransferase 2B7 (accession number P16662) related to heroin liver metabolism, with annotated sequence, quality q-value, the total number (#) of protein groups identified with the peptide, number (#) of peptide spectral matches (PSMs), peptide position in the protein, number (#) of missed cleavages of the identified peptide. * The peptide is also identified in putative inactive carboxylesterase 4 (CES1P1, accession number Q9UKY3).

| Annotated Sequence | Qvality q-value | # Protein Groups | # PSMs | Identified in Protein  accesion number [position in protein] | # Missed Cleavages |
| --- | --- | --- | --- | --- | --- |
| *[R].AISESGVALTSVLVK.[K] | 5.45707E-05 | 2 | 3 | Q9UKY3 [244-258];  P23141 [243-257] | 0 |
| [R].FTPPQPAEPWSFVK.[N] | 5.45707E-05 | 1 | 2 | P23141 [65-78] | 0 |
| [K].FVSLEGFAQPVAIFLGIPFAKPPLGPLR.[F] | 5.45707E-05 | 1 | 3 | P23141 [37-64] | 0 |
| [R].GNWGHLDQVAALR.[W] | 5.45707E-05 | 1 | 7 | P23141 [187-199] | 0 |
| [K].TVIGDHGDELFSVFGAPFLK.[E] | 5.45707E-05 | 1 | 3 | P23141 [463-482] | 0 |
| [K].MVMKFWANFAR.[N] | 0.000239472 | 1 | 1 | P23141 [495-505] | 1 |
| [K].TAMSLLWK.[S] | 0.000239472 | 1 | 2 | P23141 [377-384] | 0 |
| [K].EGYLQIGANTQAAQK.[L] | 0.00130768 | 1 | 1 | P23141 [524-538] | 0 |
| [K].FWANFAR.[N] | 0.00525625 | 1 | 1 | P23141 [499-505] | 0 |
| [R].APVYFYEFQHQPSWLK.[N] | 5.45707E-05 | 1 | 2 | O00748 [431-446] | 0 |
| [K].GANAGVQTFLGIPFAKPPLGPLR.[F] | 5.45707E-05 | 1 | 3 | O00748 [50-72] | 0 |
| [K].ADHGDELPFVFR.[S] | 0.000239472 | 1 | 3 | O00748 [455-466] | 0 |
| [K].ALPQKIQELEEPEER.[H] | 0.0048795 | 1 | 1 | O00748 [541-555] | 1 |
| [K].ADVWLIR.[N] | 1.5E-3 | 1 | 2 | P16662 [253-259] | 0 |

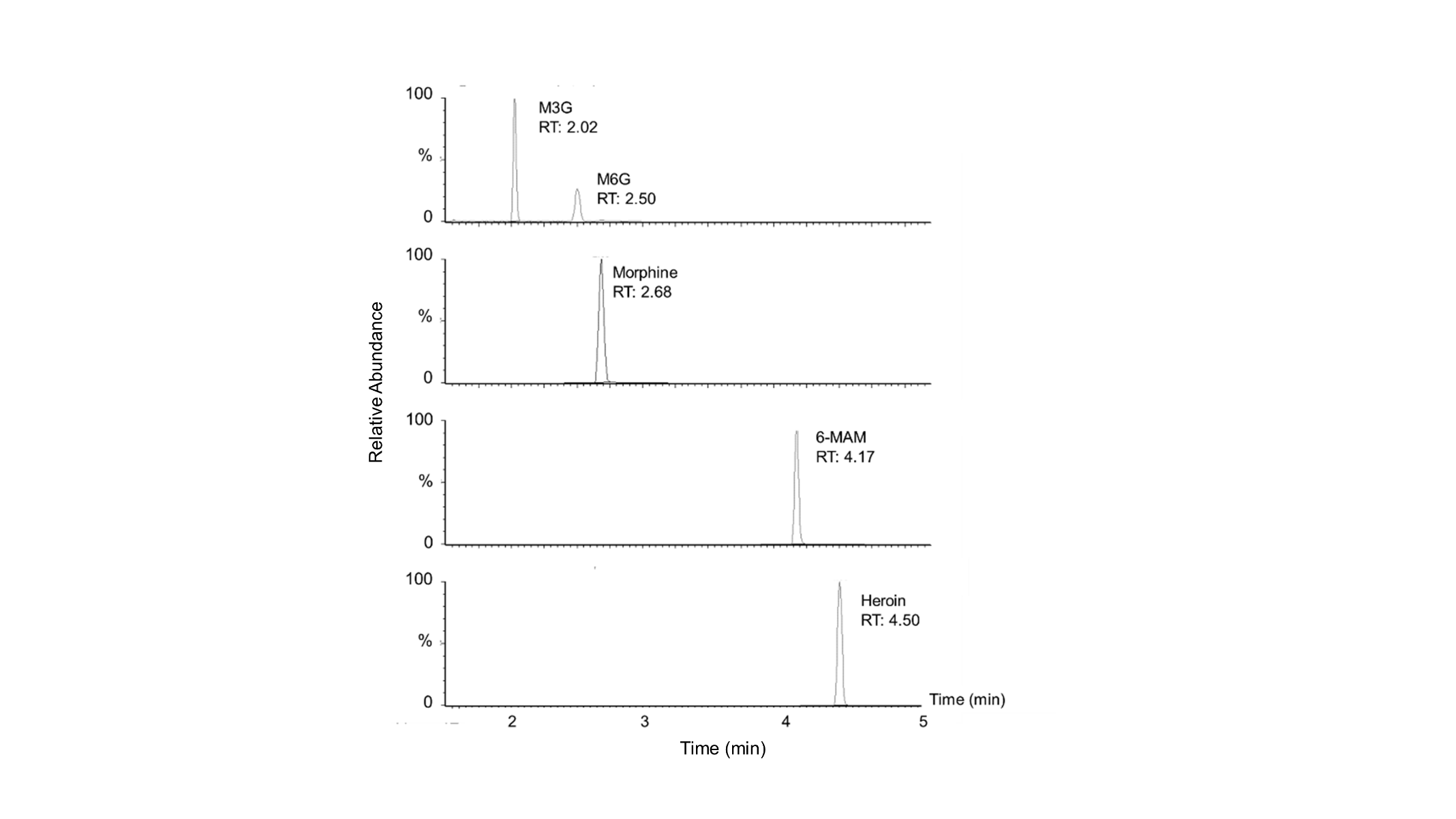

**Figure S-5.** Chromatogram showing the separation of the analytes M3G, M6G, morphine, 6-MAM and heroin with the use of UHPLC-MS.

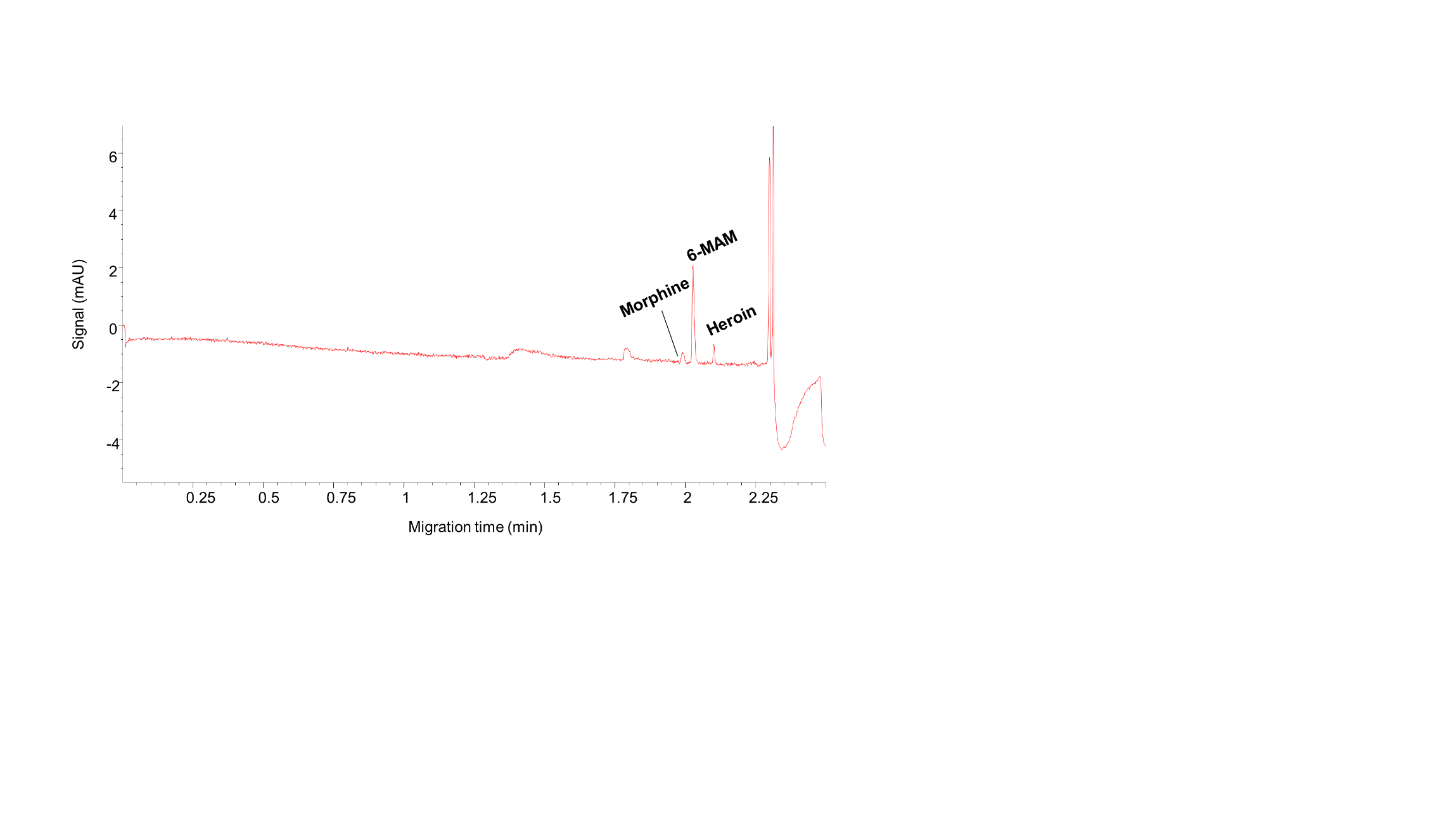

**Figure S-6.** Electropherogram of liver organoid drug phase I metabolism using Pa-EME and simple capillary electrophoresis with ultraviolet spectroscopy detection (CE-UV), after incubation with liver organoids differentiated from the iPSC cell lines AG27 (60 organoids) in 50 µM heroin for 6 hours.

**Additional experimental procedures**

**Heroin metabolism assay in microsomes and S9-fraction**

A volume of 80 µL NADPH regenerating solution (Corning Incorporated, NY, USA) and 10 µL pooled human liver microsomes (XTreme 200 pool, final concentration 0.01 mg protein/sample, XenoTech, Kansas City, KS, US) or S9-fraction (Xtreme 200 pool S9 fraction, 0.01 mg protein/sample, XenoTech) were added to Eppendorf tubes and pre-incubated at 37°C for 10 min in a water bath. The heroin metabolism was initiated by adding 10 µL 1 µM heroin (final concentration 0.1 µM), and the Eppendorf tubes were vortexed before incubation at 37°C for 0-20 minutes. The metabolism was terminated by addition of 10 µL 1.2 M formic acid followed by 10 µL of a 1.5 µM mixture of internal standards (morphine-d3, 6-MAM-d6 and heroin-d9), and the samples were immediately vortexed for 30 sec. In parallel, drug degradation control samples (without microsomes or S9-fraction) were prepared.

**Sample preparation using centrifugation**

Centrifugation was performed using a MiniSpin plus from Eppendorf (Hamburg, Germany), or a Micro Star 17R from VWR or an Allegra X-15 R from Beckman Coulter (Brea, California, USA). The samples were centrifuged at 4°C at 14 500 g for 10 min, and the supernatants were transferred to auto sampler vials. The vials were kept on ice if analyzed the same day as the experiment, or at -80°C until analysis.

**Capillary electrophoresis with ultraviolet spectroscopy detection**

The capillary electrophoresis (CE) separations with data handling were carried out using a 7100 CE instrument equipped with an on-column diode-array detector and a CE Chemstation software from Agilent Technologies (Waldbronn, Germany). Separations were performed using fused-silica capillaries from PolymicroTechnologies (Phoenix, AZ, USA), with L_tot_ = 59 cm and L_eff_ = 51 cm, 75 μm ID and 360 μm outer diameter (OD). The background electrolyte (BGE) consisted of 30 mM ammonium formate adjusted to pH 8 using a 1 M solution of NaOH. Initial conditioning of a new capillary was carried out at a pressure of 1000 mbar with 1 M NaOH (20 min), water (5 min) and BGE (20 min and for 5 min at +15 kV). Daily conditioning of the capillary was performed (1000 mbar) with 0.1 M NaOH (5 min), water (2 min) and BGE (10 min and for 5 min at 30 kV). Before each injection, the capillary was rinsed at a pressure of 1000 mbar with 0.1 M NaOH and water (0.5 min each), followed by the BGE (3 min). Hydrodynamic injections were performed at 50 mbar: starting with diluted BGE (9 +1, water + BGE) for 3 s, followed by the sample injection (15 s, equivalent to around 107 nL) and the BGE (3 s). Separations and measurements were performed with an applied potential of +30 kV (25°C) and at an UV-absorbance of 214 nm.
